# Zebrafish *carbohydrate sulfotransferase 6* (*chst6*) mutants provide a preclinical model for macular corneal dystrophy

**DOI:** 10.1101/2024.01.24.577150

**Authors:** Esra Ersoz-Gulseven, Merve Basol, Helin Özaktaş, Sibel Kalyoncu, Canan Asli Utine, Gulcin Cakan-Akdogan

## Abstract

Macular corneal dystrophy (MCD) is a rare congenital disease caused by mutations in the *carbohydrate sulfotransferase 6* (*chst6*) gene. Patients suffer from opaque aggregates in the cornea leading to bilateral progressive vision loss by 4^th^ decade of life. Corneal transplantation is the only available treatment, which is invasive, not available to every patient and recurrence of the symptoms is common. Keratocytes in the cornea express the *chst6* gene, which encodes a golgi enzyme that is essential for sulfation of the keratan sulfate proteoglycans (KSPG). The loss of KS sulfation leads to defects in collagen fibril organization and aggregate formation in the corneal extracellular matrix. Lack of preclinical disease models is a major limitation for the development of accessible treatment strategies. Attempts to develop mouse MCD models have failed due to lack of *chst6* gene in mice and difference in proteoglycan composition of the mouse cornea. The zebrafish *chst6* gene has not been studied previously. Zebrafish cornea structure is highly similar to humans, containing high levels of keratan sulfate proteoglycans in the stroma. Here, loss of function *chst6* mutant zebrafish were generated with CRISPR/Cas9 mediated gene editing. Several *chst6* alleles were obtained, and loss of KSPG sulfation in the eye stroma was shown. Mutant zebrafish developed age-dependent, alcian blue positive, opaque accumulates in the cornea. Degeneration of corneal structure and changes in epithelial thickness were observed. The zebrafish MCD model developed here is the first *in vivo* model of the disease and opens up possibilities to develop and screen treatment strategies.

**Significance Statement:** First *in vivo* model of macular corneal dystrophy (MCD) is reported in this study. Zebrafish model developed here paves the way for modeling of other corneal dystrophies in this aquatic vertebrate which is easy to apply therapeutics and image *in vivo*. The clinical symptoms of MCD are well reproduced in the zebrafish MCD model. Moreover, the authors showed that *chst6* gene function is not restricted to cornea, and a fraction of mutant larvae have morphological defects. The mutants developed here provide a genetic model for understanding the highly complex roles of keratan sulfate proteoglycans.

## Introduction

The composition and organization of the extracellular matrix (ECM) in corneal stroma is essential to its transparency (1). The keratocytes are the main architects of this ECM. They produce proteoglycans and collagens which form a perfectly aligned lattice, that remains hydrated and transparent in homeostasis. Proteoglycans play a key role in the maintenance of corneal transparency by interacting closely with collagen bundles in the eye and regulating their sizes and alignment (2, 3). Proteoglycans are complex molecules, which diversify the structure and roles of their core proteins by the type(s), number, linkage type, and the length of the glycosaminoglycan (GAG) side chains (4). The composition of the GAG content of the same core protein can vary greatly by modification of all these factors. Sulfation of the disaccharides in the GAG chains is of essential importance, hence they are mostly named as sulfates (i.e., heparan sulfate, chondroitin/dermatan sulfate, keratan sulfate) (5). Keratan sulfates (KS) are highly sulfated glycosaminoglycans (GAGs) that contain long repeats of Galactose (β1>4) N-Acetylated Glucosamine (GalGlcNac) disaccharide repeats. They are found in keratan sulfate proteoglycans (KSPGs) such as lumican, keratocan and in some complex proteoglycans such as aggrecan that carry multiple types of GAGs (6). The importance of keratan sulfates and their sulfation status in the cornea extracellular matrix are exemplified by diseases cornea plana type 2 and macular corneal dystrophy (MCD) that are linked to mutations in *keratocan* and *carbohydrate sulfotransferase 6*, respectively (7, 8). The structure and composition of cornea is highly conserved; hence the zebrafish eye is a promising model for studying proteoglycan related corneal diseases.

Macular corneal dystrophy is characterized by opaque aggregates that begin to accumulate in the cornea in early teen years and gradually increase to finally cause bilateral vision loss by the 4^th^ decade of life which is treated with corneal transplantation (9). Although MCD is a rare disease with autosomal recessive transmission, its incidence is higher in populations with high consanguinity including South India, Turkey, Iceland (10-12). 25% of corneal transplants are performed to corneal dystrophy patients and MCD is the most common corneal dystrophy in Turkey (13). Access to corneal donors is a limitation even in developed countries while a significant proportion of the world population has major problems in accessing this treatment (14). Moreover, the symptoms recur after transplantation as the keratocytes of the patient invade the transplanted cornea (15, 16). Lack of preclinical *in vivo* models is a limitation for development and testing of treatment strategies.

MCD is caused by mutations in *carbohydrate sulfotransferase 6* (*chst6*) gene which encodes for a N-acetylglucosamine-6-O-sulfotransferase (GlcNAc6-ST), major enzyme for sulfation of keratan sulfate proteoglycans (KSPGs) (MIM #217800) (17). Lack of sulfated corneal keratan sulfate (cKS) in corneal stroma is common for all MCD patients, whereas some also lack cKS in keratocytes and serum (18, 19). Corneal KSPGs are in contact with collagen fibrils, and abnormalities in bundle sizes and organization of collagen fibrils are detected in MCD patient samples (2, 20). Negative charge of sulfate groups is proposed to be important for the hydration of corneal extracellular matrix (ECM) and collagen bundle size control by physical force (3). Experimental demonstration of the disease mechanisms is yet to be done, since there are no *in vivo* disease models available. Although an *in vitro* model via inhibition of sulfotransferases in cultured goat primary corneal cells was proposed, lack of ECM containing collagen bundles and proteoglycans that are tightly organized is a major limitation of this 2D model (21). In humans, highly similar *chst5* and *chst6* genes encode for intestinal and corneal specific GlcNAc6-ST enzymes, respectively. On the other hand, mice have only *chst5* as ortholog of these two genes. It was not possible to mimic MCD in mice although null mutants of *chst5* were generated (22). Which may be explained by the fact that mouse cornea contains predominantly chondroitin sulfate proteoglycans (CSPGs), whereas 80% of human corneal proteoglycans are KSPGs (22). Zebrafish has a *chst6* gene and is in a unique position to model MCD, since the structure of cornea is well conserved. Zebrafish corneal stroma is composed of keratocytes and precisely organized collagen lattices (23, 24). Importantly, 80% of the proteoglycans in zebrafish cornea are KSPGs (25). Moreover, in zebrafish the main corneal KSPGs *lumican* and *keratocan* are conserved and stromal localization of cKS is detected as of larval stages (Zhao et al., 2006, Yeh et al., 2008, 2010). No previous study reported expression and function of *chst6* in zebrafish, while expression of *chst6* in embryonic zebrafish reported in ZFIN (26). Here, it was aimed to generate loss of function *chst6* mutant zebrafish to generate a preclinical MCD model.

## Results

### *Carbohydrate sulfotransferase 6* expression during development

Whole mount *in situ* hybridization revealed that *chst6* expression in head structures and the trunk 24 and 30 hours post fertilization (hpf). At 60 - 72 hpf, the stages corneal layers are forming in zebrafish eye, *chst6* expression becomes enhanced in the anterior tissues including the brain, eye, and jaw. Close-up images show strong *chst6* expression in the eye and sections of the stained sample shows expression in the retina and cornea of the larval eye (Fig. 1).

**Figure 1.**
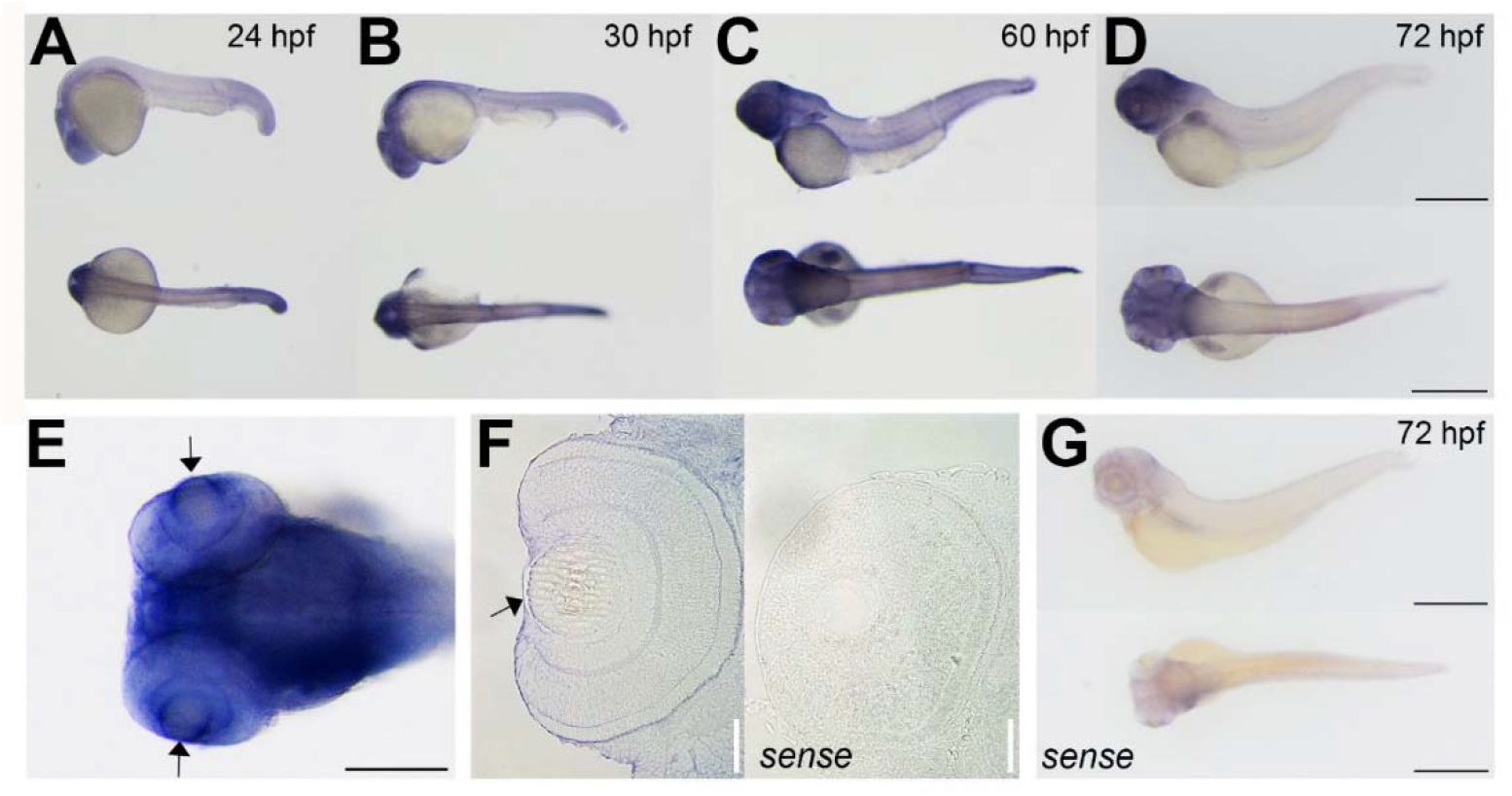
The expression pattern of *chst6* in zebrafish embryos. Whole mount in situ hybridization revealed expression of *chst6* in the body **A)** 24 hpf, **B)** 30 hpf, **C)** 60 hpf, and **D)** 72 hpf zebrafish. **E)** Close-up image of the 72 hpf larval head shows strong expression in the brain and the eye. **F)** Section of larvae post in-situ hybridization with *chst6* antisense and sense (negative control) probes, shows *chst6* expression in the cornea (arrow). **G)** Representative images of 72 hpf larvae stained with sense (negative control) *chst6* sense probe. Scale bar: 100 μm.

### Zebrafish *chst6* mutants were generated via CRISPR/Cas9 mediated gene editing

Large number of mutations distributed to the entire coding sequence of *chst6* were reported in MCD patients, without any definite hotspot (27). To choose the best target for CAS9, the structure of the CHST6 protein (UNIPROT ID: Q9GZX3) was homology-modelled via SWISS-MODEL by using the crystal structure of *Mycobacterium avium* sulfotransferase protein (PDB ID: 2Z6V) as a template. The protein has a transmembrane domain (6 - 26) that ensures golgi localization and a lumenal sulfotransferase domain (27-395) (Fig. 2A). To predict the enzyme active site, the homology-modelled structure was analyzed by PyMOL. It was found that the 1^st^ and 2^nd^ 3’-phospho-5’-adenylyl sulfate (PAPS) binding sites form a sulfate passageway in the folded enzyme, while the arginine in 1^st^ PAPS binding region (WRSGSSF) was found to bind the SO_4_ donor suggesting it is the enzyme active site (Fig. 2B). This site is fully conserved among several vertebrate species including the zebrafish, and all residues but the first W are found to be mutated in patients (27). Based on this knowledge, deletion or inactivation of 1^st^ PAPS site was chosen as a strategy to generate *chst6* mutants. Guide RNAs (gRNAs) were designed with CRISPRscan to induce 1) induction of a double strand break (DSB) before the 1^st^ PAPS binding site, 2) induction of a 50 bp deletion spanning the 1^st^ PAPS site, 3) induction of an early DSB, 4) deletion of the cds with 2 gRNAs (Fig. 2A, Table 1) (28).

**Table 1.**
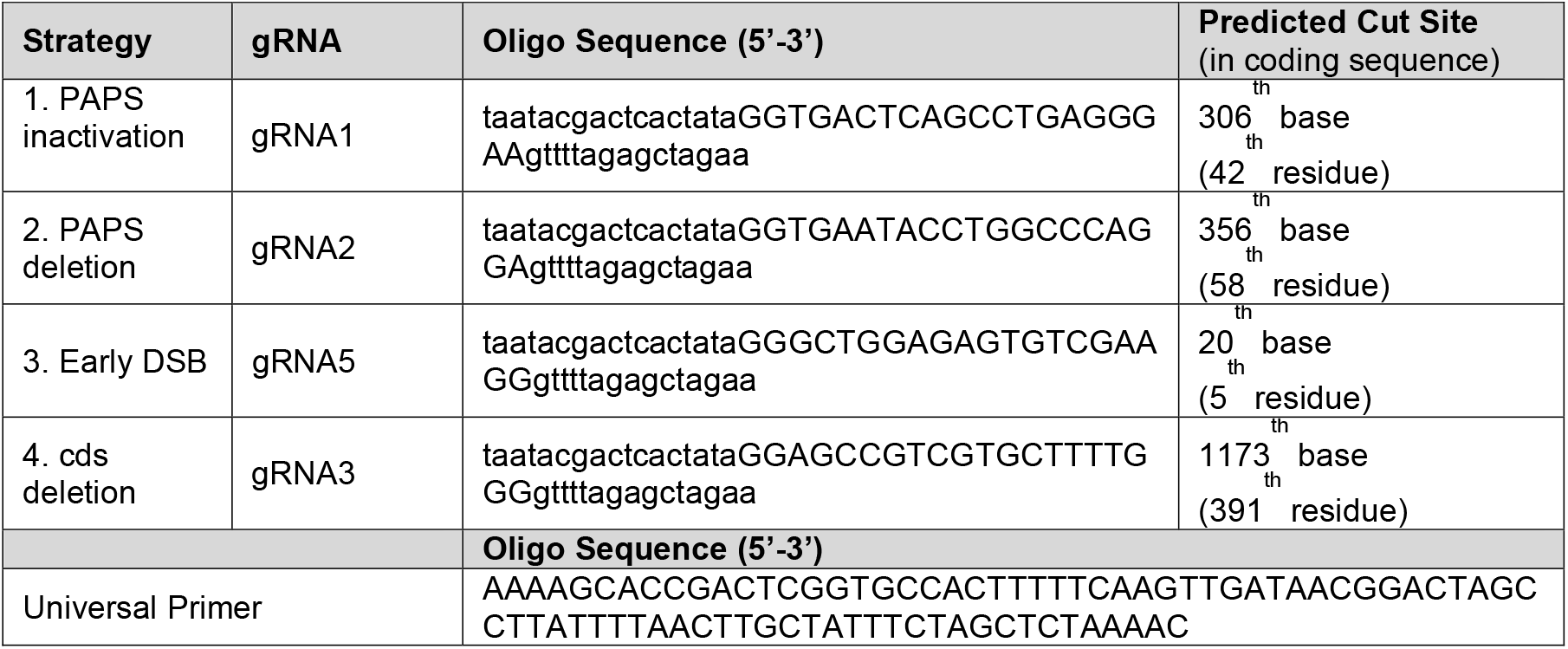
Gene specific and universal oligos used to generate guide RNAs and respective predicted cut sites. Gene specific sequences are shown uppercase in gRNA specific oligos.

**Figure 2.**
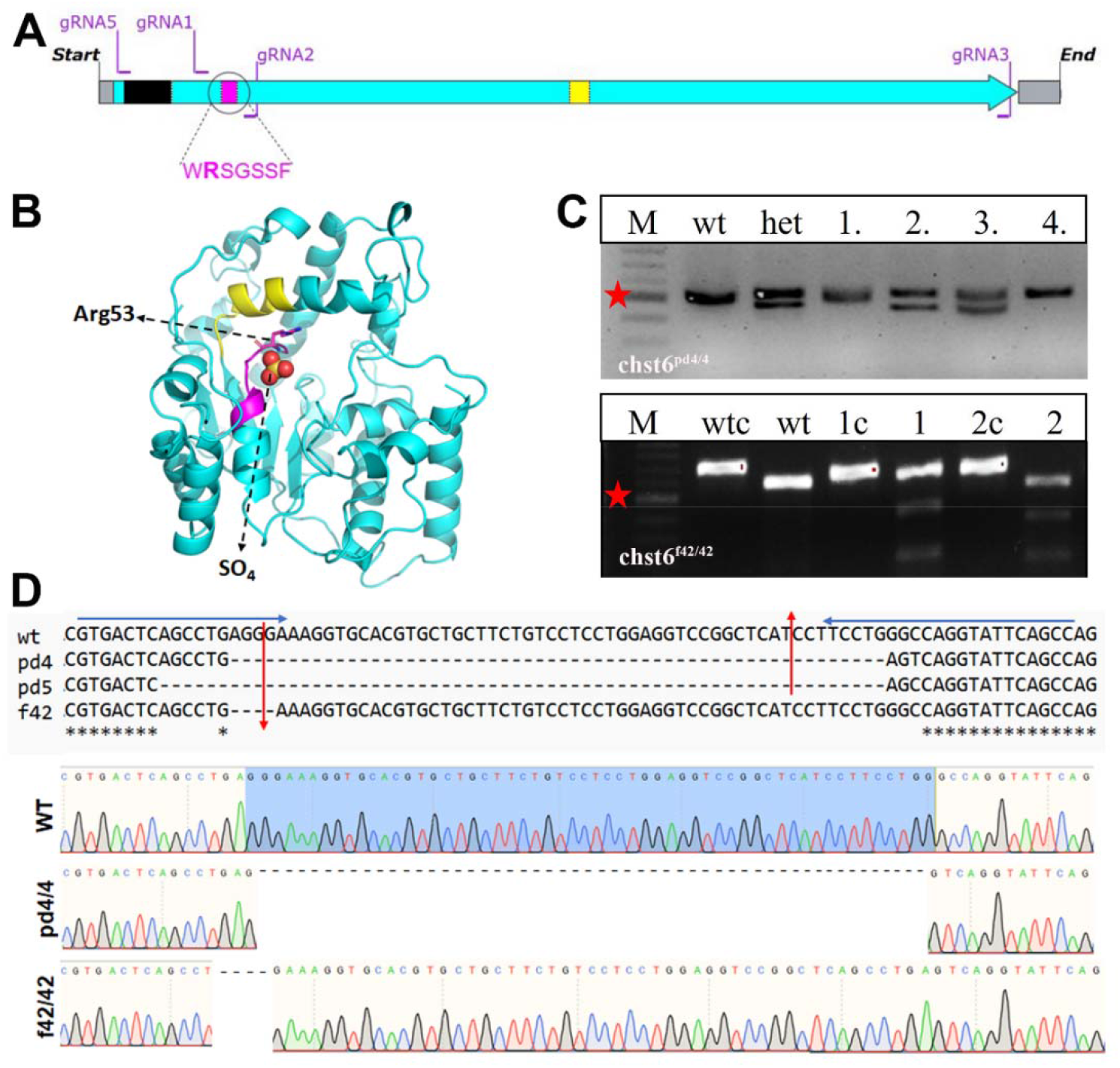
*chst6* mutagenesis strategy: **A)** Graphical representation of *chst6* gene coding sequence (cds). Sequences coding carbohydrate sulfotransferase domain (turquiose), 1^st^ PAPS (magenta) and 2^nd^ PAPS (yellow) and transmembrane site (black) were colored, and gRNA binding sites were indicated. **B)** Model of zebrafish CHST6 sulfotransferase domain indicating localization of 1^st^ PAPS (a.a. 49-55, colored magenta), and 2^nd^ PAPS (a.a. 202-210, colored yellow) and SO_4_ interaction with Arg53. **C)** Representative agarose gel images of genotyping deletion and frameshift mutants, red asterisk shows 500 bp. **D)** Multiple sequence alignment of wildtype *chst6* gene and homozygous mutants, pink: 1^st^ PAPS site, blue arrow: gRNAs, red arrows: predicted cut positions.

Although the highest scoring gRNA (gRNA5) targeted the 5^th^ residue, no mutant alleles were obtained with this gRNA (Table S1). Use of gRNA5 together with gRNA3 resulted in mosaic fish carrying 943 bp deletions, and deletion started at 284^th^ bp, 250 bp downstream of the predicted gRNA5 cut site (Fig. S1). However, only heterozygous fish were obtained with this large deletion and no mutant line was established.

gRNA1 and gRNA2, that are predicted to target base pairs coding for 42. and 58. residues, respectively were designed for deleting the 1^st^ PAPS region. Resulted deletion mutations were identified by gel mobility shift of a PCR product that amplified 522 bp genomic DNA covering the target sites, and gRNA1 induced indel was detected with a T7EI mismatch assay (Fig. 2C). Use of gRNA1 alone and gRNA1+ gRNA2 both resulted in successful mutagenesis of the targeted regions, leading to generation of 13 and 6 founder fish, respectively (Table S1). The sequences of selected alleles are shown in Fig. 2D and Fig. S2. Lethality was observed in the larval-juvenile stages; however, it was less pronounced in the next generations and surviving homozygous fish were sufficient to establish lines (Table S2). Mutant larvae displayed morphological defects at low penetrance, which was rescued by wt mRNA injection (Fig. S3). Only the larvae with normal morphology were raised to adulthood and in the next generations penetrance of body defect phenotype was much reduced (Fig. S3). All of the following experiments were conducted with at least 2 alleles and in two different generations.

### The zebrafish *chst6* mutants lost sulfated keratan sulfates

In order to test for loss of function in the mutants, antibodies raised against N-terminal epitopes of human CHST6 were used, however there was no cross-reactivity with zebrafish CHST6. Next, the 5D4 cKS antibody which recognizes the fully sulfated GlcNac-Gal epitopes of keratan sulfate was used (23). ELISA assay was used to quantify cKS in the whole-body larval lysates. The total amount of cKS per larva was 65.7 pg in WT larvae, while it was 15.4, 17.7 and 25.9 pg per *chst6*^*pd4/pd4*^, *chst6*^*pd5/pd5*^ and *chst6*^*f242/f42*^ larvae, respectively (Fig. 3A). Immunofluorescence staining showed that cKS signal is lost in the cornea and head of the mutants (Fig. 3C-C’, Fig. S4). Injection of wt *chst6* mRNA to zygotes restored cKS expression (Fig. 3D-D’).

**Figure 3:**
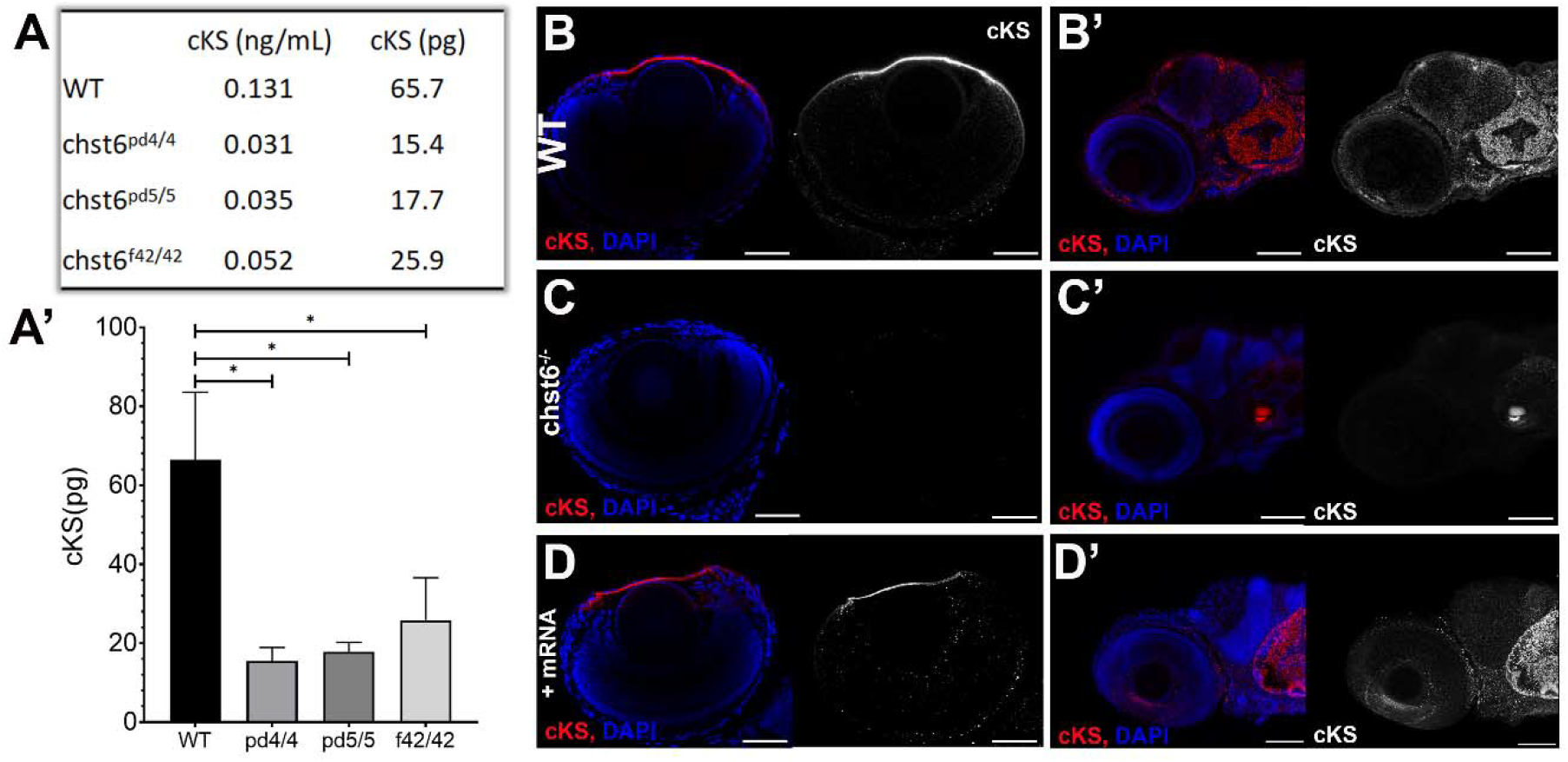
Zebrafish mutants have defective keratan sulfate sulfation. **A)** Table **A’)** graph showing total cKS amount in wildtype and homozygous larvae whole body extracts. Representative images of cKS antibody staining: **B)** eye and **B’)** head of wt larva and **C)** eye and **C’)** head of chst6^pd4/4^ mutant larva, **D)** eye and **D’)** head of homozygous mutant larvae that was rescued with *chst6* wt mRNA. Scale bar: 60 μm.

### Macular corneal dystrophy symptoms were reproduced in zebrafish *chst6* mutants

Opaque aggregates were detected in mutants (Fig. 4). This phenotype was seen earliest at 8 months post fertilization (mpf), the incidence of MCD phenotype increased over time and most homozygous mutants developed opaque deposits between the ages of 14 mpf - 20 mpf (Fig. 4A-E, Fig. S5). These aggregates impaired vision which was demonstrated by reduced response of mutant fish to food, dispersed on the surface of water (Fig. 4F, Fig. S5B, Supplementary videos).

**Figure 4:**
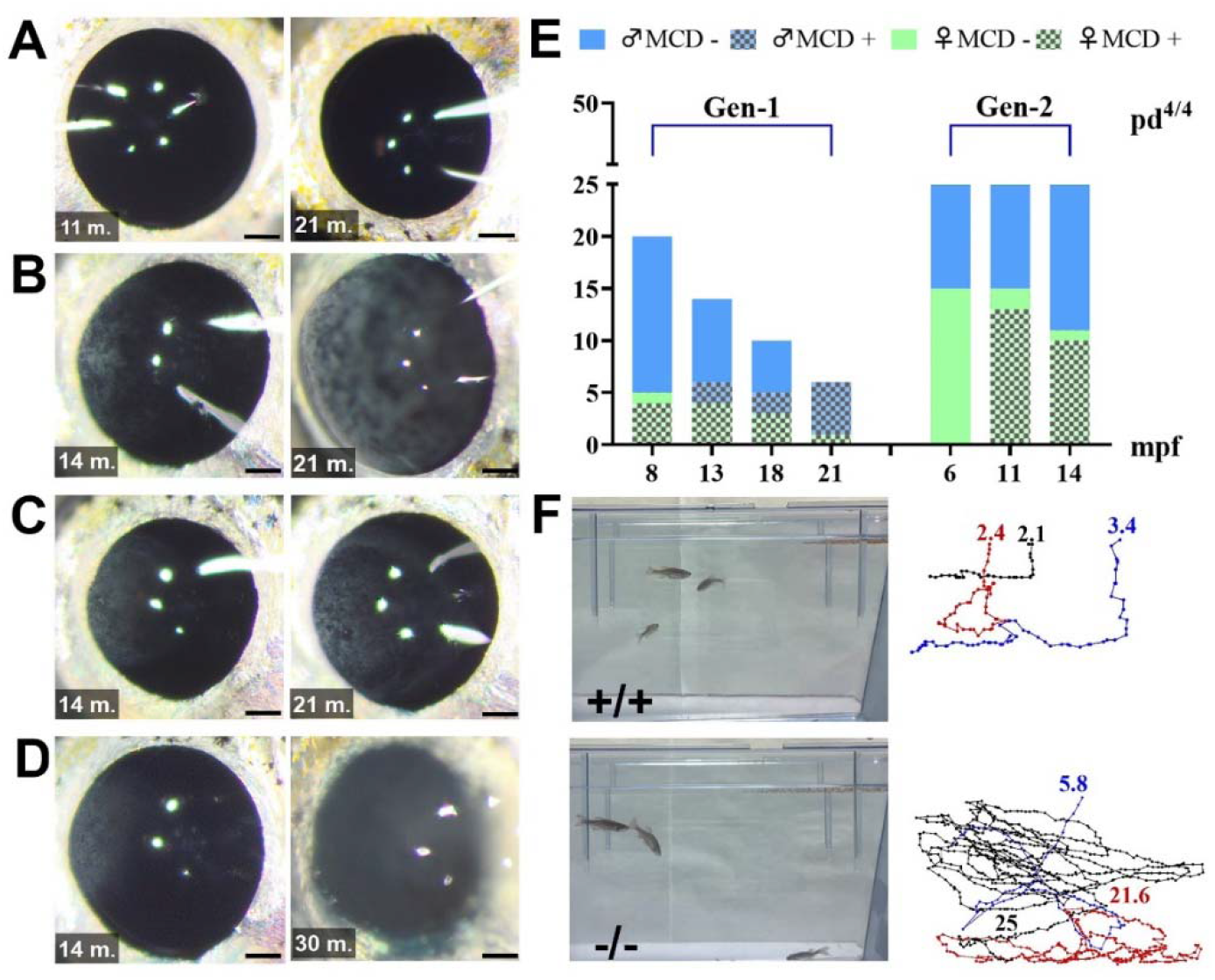
Opaque aggregates in the cornea and impaired vision. Macroscopic analysis of **A)** WT, **B)** *chst6*^*pdpd/4*^, **C)** *chst6*^*pd5/pd5*^, and **D)** *chst6*^*f42/f42*^ alleles. Eyes of (Left) young (11-14 mpf) adults of generation 2, (right) old (21-30 mpf) adults of generation 1 are shown. Scale bar: 300 μm. **E)** Incidence of opaque corneal aggregates homozygous mutants of two generations over time. Green bar: females, blue bar: males, dashed bars indicate occurrence of opaque aggregates. **F)** The Vision test was performed with WT (+/+) and homozygous chst6 mutants (-/-). Swim path and duration of fish were tracked. Track was ended if the fish reached food and time stamp is displayed at the end of each track. The mutants swam for 21.6 and 25 seconds without noticing the food.

Next, paraffin sections of adult fish were stained with alcian blue and periodic acid schiff. In the cornea of wt zebrafish the epithelium was stained strongly positive with alcian blue while some positive foci were detected in the stroma (Fig. 5A-A’). Stroma of mutants were compact and wt and blue deposits were detected (Fig. 5B-B’). Some mutants had degenerated posterior stroma/descement membrane and blue deposits on this level (Fig. 5C-C’). The stroma was thinner in most mutants and some lost stroma almost completely (Fig. S6). One very old mutant (30 mpf) carrying the *chst6*^*f242/f42*^ allele had atypical cornea structure, which has an overgrown and irregular epithelium and very thick cornea (Fig. 4D, Fig. S6G).

**Figure 5:**
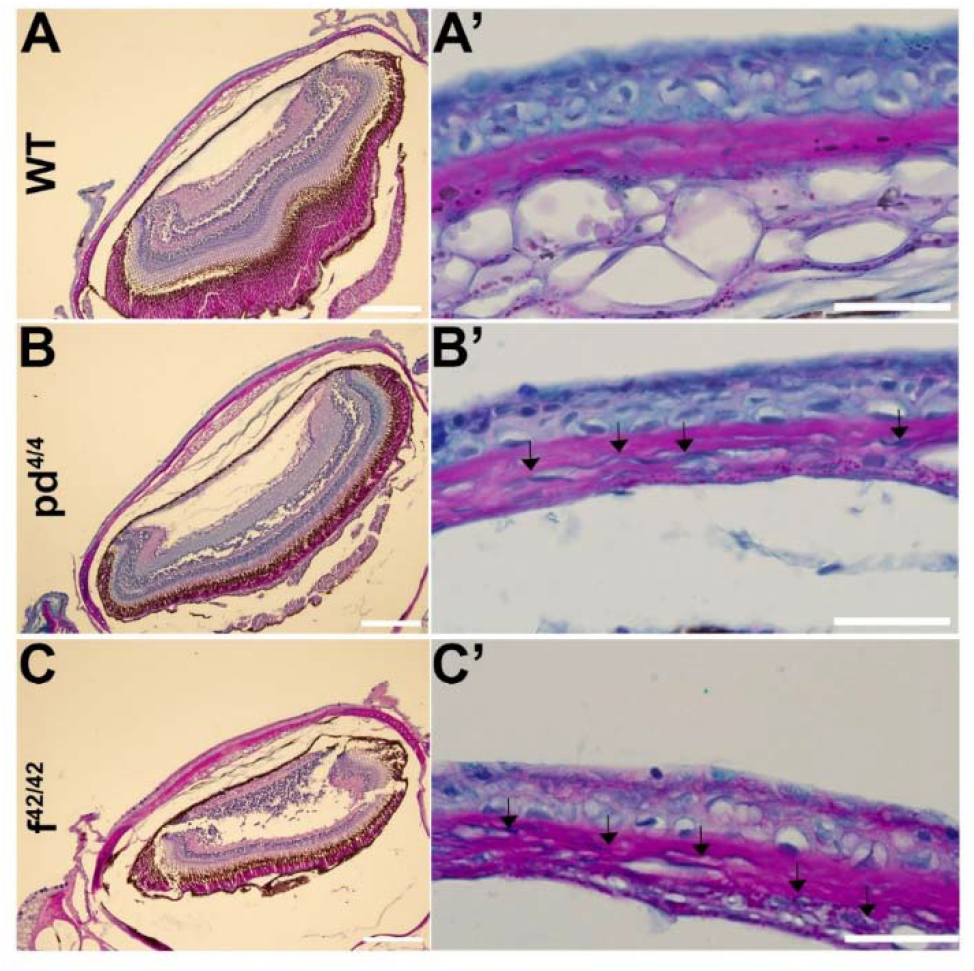
Zebrafish with MCD have alcian blue positive aggregates in the cornea. Alcian blue/PAS-stained eye sections of **A)** WT, **B)** *chst6*^*pd4/pd4*^ and **C)** *chst6*^*f42/42*^ show overall eye structure. Cornea close-up images of **A’)** WT, **B’)** *chst6*^*pd4/pd4*^ and **C’)** *chst6*^*f42/42*^ adult zebrafish show epithelial and stromal structure. Black arrows indicate alcian blue positive aggregates. Collagen in stroma is stained pink with PAS. Scale bar: (A-C): 200 μm, (A’-C’) 30μm.

**Figure 6:**
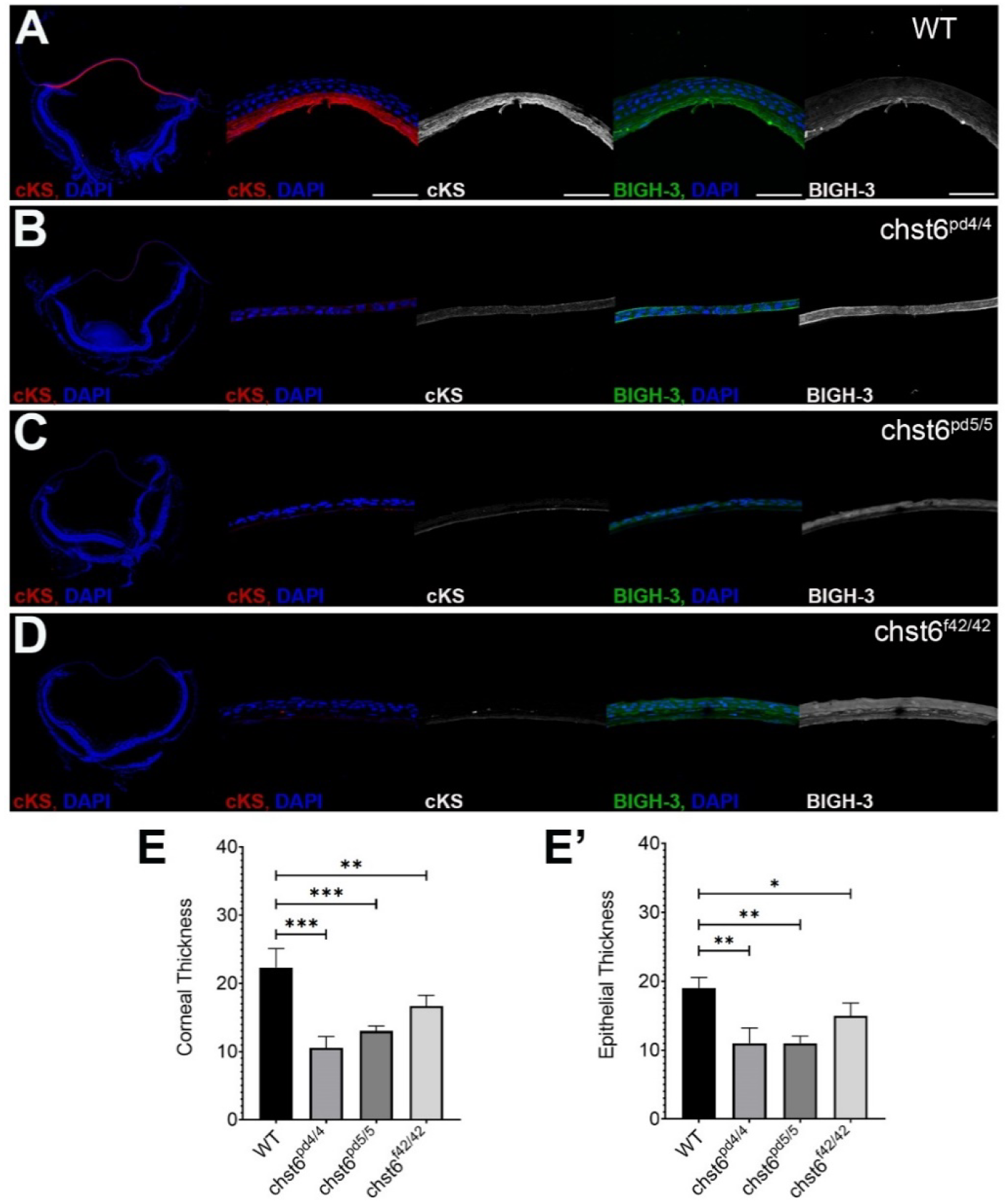
cKS was depleted in cornea of homozygous *chst6* mutants. Eye sections stained with anti-cKS(red), anti-BIGH3 (green) and DAPI (blue) are shown. **A)** wildtype **B)** *chst6*^*pd4/pd4*^ and **C)** *chst6*^*pd5/pd5*^ and, **D)** *chst6*^*f42/42*^ mutant. Average thickness of **E)** cornea, **E’)** corneal epithelium (n=5). Scale bar: 80 μm.

Another characteristic of MCD cornea is loss of cKS in the corneal stroma. While cKS was present in the corneal stroma of wild type zebrafish, it was completely lost in all homozygous mutants (Fig. 5). TGF-beta induced protein Ig-H3 (BIGH3) is an epithelial marker in human cornea. Double staining of cKS and BIGH3 in wild-type fish showed that BIGH3 was present in the epithelium and at low levels in the stroma of zebrafish adult eye. In the mutants BIGH3 expression in the corneal epithelium was not affected, whereas it was lost in the stroma of *chst6*^*pd4/4*^, *chst6*^*pd5/5*^, and decreased in the stroma of *chst6*^*f42/42*^ homozygous mutants (Fig. 5 A-D). Moreover, the thickness of the cornea and the cornea epithelium was decreased in all mutants (Fig. 5 E-E’). These data showed that the histopathological aspects of MCD phenotypes were well reproduced in the zebrafish mutants reported here.

## Discussion

Macular corneal dystrophy patients were reported to have different mutations causing enzyme loss-of-function. Conservation among species and frequency of mutagenesis for each residue was reported by Zhang et. al. Density of the mutations was highest between 46-77. aa, with 25 out 32 residues mutated in multiple patients, and 23 of these mutated residues are identical in zebrafish CHST6 protein (27). By analyzing the homology model of CHST6 protein, we showed that R53 located in 1^st^ PAPS domain binds the sulfate donor, hence is most likely to be the enzyme active site. We also confirmed the accuracy of our modelled structure with later published AlphaFold predicted model (AF-Q9GZX3-F1). Inactivation of 1^st^ PAPS by introduction of an indel via gRNA1 or a deletion via gRNA1 and gRNA2 targeted CAS9 both resulted in successful mutagenesis and generation of loss-of-function mutants. Our attempts to generate mutants with deletion of the entire coding sequence was not successful. Even though the designed gRNA5 had highest score according to specific algorithms that was developed based on on empiric data, the deletion did not occur at the 5^th^ a.a. as predicted. Instead, a fairly large deletion that starts from 89^th^ a.a. was generated, suggesting that either the DSB was not possible at this locus or it was repaired. The dynamics of Cas9 catalytic activity has been shown to vary at different target sites and chromosome opennes at the target is proposed to be a factor, which may be the case here (29, 30). The fact that only heterozygote zebrafish was obtained with the large deletion, may indicate a vital role for integral *chst6* genomic region for survival.

Loss-of-function was proven by loss of enzyme activiy in the mutants. 5D4 antibody was used in this study to assess enzyme function in *chst6* mutants. Quantification with ELISA showed that the amount of cKS was decreased strongly in whole body of double mutant larvae but did not completely dissappear. In the zebrafish genome *chst2* and *chst6* are the only genes encoding keratan sulfate GlcNac sulfotransferases. Although neither of these two genes were studied in depth, *chst2* expression was reported in the central nervous system (31). The remaining cKS may be due to the activity of *chst2a* and/or *chst2b* genes. Immunofluorescence staining showed that corneal localized KS signal was lost in mutant larvae, and this was rescued when wildtype mRNA was injected to 1-cell stage embryos. Similarly, in adult zebrafish stromal cKS signal was completely lost in homozygous *chst6* mutants.

In humans, CHST6 is predominantly present in cornea whereas CHST5 takes on the same function in the intestine. We showed that *chst6* expression is initially ubiquitous in 24 - 30 hpf embryos whereas it becomes restricted to the head region by 72 hpf. Expression in the eye and cornea was detected at 72 hpf, which corresponds to a stage of corneal layer specification in zebrafish eye (23). In parallel to the expression of *chst6* in the head and cornea, sulfated keratan sulfate was detected in these tissues of 4 dpf wt zebrafish and lost in the chst6 mutants. These findings suggest that zebrafish *chst6* gene gained additional functions outside of cornea. Supporting this, a small portion of mutant embryos were malformed and did not survive (Fig. S4). Low penetrance of such phenotype indicates a complex regulatory mechanism which is beyond the scope of this manuscript and will be the focus of future investigations. Here, the otherwise healthy-looking mutant larvae were raised, and corneal stroma was investigated in the context of MCD disease.

Some MCD patients also display loss of cKS in the serum indicating a systemic effect which is not yet understood completely (32). Indeed, there are three variants of MCD in humans, characterized by immunophenotype. Type 1 patients have no detectable keratan sulfate in either the serum or cornea. In type 1A patients, keratan sulfate is absent in the serum but stroma shows immunoreactivity to keratan sulfate antibodies, whereas in type 2 normal amounts of keratan sulfate is detected in the serum and stroma (33). The zebrafish mutants have no cKS in the cornea but a low cKS is present in the whole body at the larval stages, serum of the fish was not tested for cKS due to very low volume of the blood.

In the zebrafish MCD model described here, different mutant alleles were characterized through different generations and consistent results were obtained regardless of the allele. Formation of opaque aggregates in the eye occurred in adulthood as early as 8 months post fertilization, incidence of opacity increases as the fish became 14 months or older. cKS signal in the corneal stroma was completely lost, however alcian blue deposits were detected in both the epithelium and stroma of the mutant zebrafish. Moreover, the thickness of cornea was reduced in mutants, more pronounced thinning was detected in the epithelium. These structural changes in zebrafish cornea reflects the MCD patient symptoms as well. In humans, the corneal deposits are not present at birth but starts to be seen in the 1^st^ decade or around adolescence. The grayish white punctate opacities merge into larger areas overtime, causing the entire corneal stroma to become cloudy. The corneal stroma between the deposits is also hazy, so that the vision is disturbed and necessitates corneal transplantation around 3^rd^ – 4^th^ decades (34).

Few aspects of MCD in zebrafish differs from that of human. In zebrafish MCD, females developed opacities earlier than males, however no gender predilection was reported in humans. BIGH3 was found to be ecpressed at low levels in the corneal stroma of wt adult zebrafish, whereas in humans it is solely epithelial (23). Interestingly, BIGH3 expression was lost or decreased in the stroma of *chst6* homozygous mutants. In humans, mutations in the BIGH3 gene encoding for keratoepithelin protein have been described in different corneal dystrophies such as granular corneal dystrophy, lattice corneal dystrophy, and their different clinical subtypes, but not the MCD (35). Yet, a link between KSPGs and TGF-beta is proposed by several studies. Induction of TGFB by lumican was shown in a joint fibrosis model (36). Moreover, an interaction between lumican and TGF-beta pathway was reported in tumor progression and STRING database shows a direct interaction between lumican and TGFBI (37, 38). Our finding in zebrafish cornea also indicates a close relationship between KSPGs and TGFBI distribution in the ECM of zebrafish, which is a concept to be explored further.

In conclusion an *in vivo* zebrafish model of MCD was developed by CRISPR/Cas9 mediated mutagenesis of *chst6* gene. To our knowledge this is the first report of MCD animal model and first study showing that zebrafish cornea can be used to model corneal dystrophy. Patient symptoms and structural changes in the MCD cornea were well reproduced in the zebrafish model which paves the way to use this as a preclinical test model. Future studies will focus on development and testing of treatment approaches.

## Materials and Methods

### Zebrafish Maintenance

Zebrafish were reared under standard conditions in Izmir Biomedicine and Genome Center Zebrafish Facility. Wild-type AB strain was used to generate mutant lines. All experiments were approved by the IBG local ethics committee (IBG-HADYEK) with protocols number 202323 and 21/2019. Homozygous mutants were outcrossed to wildtype to clean the background and avoid inbreeding.

#### *In situ* Hybridization and Rescue Experiment

Whole mount *in situ* hybridization was done as published before (39). Coding sequence of *chst6* was amplified with forward 5’ATCGTGCTCGAGATGGCAAA, reverse 5’AATGCACAAATGCCCCAGAA3’ primers and cloned into pGEMT vector for full length sense probe. A full-length antisense probe was synthesized from this plasmid. A short probe was synthesized from PCR product generated with Forward 5’ATGCTGCGCTGGAGAGTGT3’ and Reverse 5’TAATACGACTCACTATAGGGAATGCACAAAT-GCCCCAGAA3’ primers. Probes were transcribed with TranscriptAid T7 High Yield Transcription kit (Thermo, KO441) using DIG-Label UTP (Roche,11127073910).

For the rescue experiment, the *chst6* coding sequence was subcloned into PCS2, linearized with NotI and mRNA was synthesized with mMESSAGE mMACHINE™ SP6 Transcription Kit (Invitrogen, AM1340). mRNA (400 ng/μl) was injected to zygotes.

### Generation of Mutants with CRISPR/Cas9

Guide RNA (gRNA) sequences were designed with the CRISPRscan web tool, and gRNAs were synthesized according to published protocols (28, 40). Cas9 mRNA was synthesized from plasmids (28, 41). pCS2-nCas9n was a gift from Wenbiao Chen (Addgene plasmid #47929; http://n2t.net/addgene:47929; RRID: Addgene_47929). pCS2-nCas9n-nanos 3’UTR was a gift from Antonio Giraldez (Addgene plasmid # 62542; http://n2t.net/addgene:62542; RRID: Addgene_62542). 250 ng/μl of Cas9 mRNA and 50 ng/μl of gRNA(s) were mixed with phenol red and ultrapure water in 10 μl volume. The mixture injected into the one-cell stage of wildtype AB embryos.

### Quantification of cKS with ELISA

5 dpf larvae were collected and snap-freeze with liquid nitrogen. At least 50 larvae per assay were used. Frozen larval pellet was crushed manually with a micro pestle, in 50 μl PBS that contains 1μl protease inhibitor cocktail (Abcam, ab201111). Tissue debris was removed with centrifugation at 12000 rpm, 4 °C for 5 minutes. Total protein was calculated by the BCA assay (Thermo, 23227). 50 μl of standard, blank, and samples were added into the antibody-coated wells. ELISA was performed using the kit manufacturer’s (Mybiosource MBS288502) protocol. The standard graph was plotted with the non-linear 4-parameter sigmoidal curve method, and the concentration of samples was calculated using Excel software.

### Whole mount Immunofluorescence Staining

The larvae were fixed with 4% PFA for overnight at 4 °C, then dehydrated with PBS/Methanol series, and kept in 100% Methanol at -20 °C for up to 24 hours. The larvae were transferred to PBS and washed with 1X PDT (PBS, 0.1% Tween-20, 0.3% Triton-X, 1% DMSO), 2 times for 30 min each. Then samples were incubated for 1 hour at 37 °C with 10 μg/μl proteinase K or 0.5 U/ml chondrotinase (SIGMA, C3667-5UN) to ensure penetration. Samples were incubated with blocking buffer (PBS, 0.1% Tween-20,10% Normal Goat Serum, 2% BSA) for 2 hours at room temperature. The larvae were incubated with the primary antibody, 5D4 Anti-Keratan Sulfate (1:200, Millipore MAB2022) for overnight at 4°C, and secondary antibody for 2 hours at room temperature. DAPI (D9542-5MG, SIGMA) was used for nuclear counterstaining. Samples were visualized with the confocal microscope (ZEISS LSM880), 25X, 40X Water and 63X objectives were used to capture Z-stacks, background subtraction and maximum Z-projection were applied with ImageJ.

### Imaging of adult eye

Adult zebrafish were anesthetized using 0.04% MESAB. The zebrafish eyes were imaged with a stereo microscope with 4X magnification, with top illumination. After examination, the fish were promptly returned to the system water. Procedure was repeated every two months.

### Vision Test with adult fish

Wildtype and mutant adult fish in their rearing tanks were kept in equal numbers, time for adjustment to environment was given to fish before food was dispersed. Video recordings were taken 1 minute before and after feeding. The second in which the feed was taken was considered to be time zero. The time it takes for the fish to reach the feed was recorded and the path of each fish to reach the feed was tracked on the video with Avidemux software.

## Acknowledgements

This study was funded by TUBITAK Grant no 219S943, EEG, and HO were supported by TUBITAK-BIDEB fellowships. Authors thank Emine Gelinci and Ecem Uzun for help with the histopathology procedures, Emine Gelinci and Meryem Ozaydın for excellent fish care. Authors thank IBG Optical Imaging and Histopathology Core facilities and IBG Zebrafish Unit.

## Acknowledgements

This project was funded by TUBITAK with grant no 219S943. Authors thank Izmir Biomedicine and Genome Center Zebrafish Facility, Histopahtology Core Facility and Optical imaging core Facility, for supporting the experiments. Authors extend gratitude to MSc Emine Gelinci and Ece Uzun for excellent help with histopathology procedures.

## References

1. J. R. Hassell, D. E. Birk, The molecular basis of corneal transparency. Exp Eye Res 91, 326–335 (2010).

2. S. Akhtar, H. M. Alkatan, O. Kirat, A. A. Khan, T. Almubrad, Collagen Fibrils and Proteoglycans of Macular Dystrophy Cornea: Ultrastructure and 3D Transmission Electron Tomography. Microsc Microanal 21, 666–679 (2015).

3. P. N. Lewis et al., Structural interactions between collagen and proteoglycans are elucidated by three-dimensional electron tomography of bovine cornea. Structure 18, 239–245 (2010).

4. J. C. Ulf Lindahl, Koji Kimata, Jeffred D. Esko, Chapter 17 Proteoglycans and Sulfated Glycosaminoglycans, Essentials of Glycobiolpgy (Cold Spring Harbor Laboratory Press, 2017).

5. D. Soares da Costa, R. L. Reis, I. Pashkuleva, Sulfation of Glycosaminoglycans and Its Implications in Human Health and Disorders. Annu Rev Biomed Eng 19, 1–26 (2017).

6. B. Caterson, J. Melrose, Keratan sulfate, a complex glycosaminoglycan with unique functional capability. Glycobiology 28, 182–206 (2018).

7. N. S. Pellegata et al., Mutations in KERA, encoding keratocan, cause cornea plana. Nat Genet 25, 91–95 (2000).

8. F. Niel et al., Truncating mutations in the carbohydrate sulfotransferase 6 gene (CHST6) result in macular corneal dystrophy. Invest Ophthalmol Vis Sci 44, 2949–2953 (2003).

9. G. K. Klintworth, C. F. Smith, Macular Corneal Dystrophy. American Journal of Pathology 89, 167–182 (1977).

10. F. Jonasson et al., Macular corneal dystrophy in Iceland. A clinical, genealogic, and immunohistochemical study of 28 patients. Ophthalmology 103, 1111–1117 (1996).

11. F. Tuncay et al., Genetic analysis of CHST6 and TGFBI in Turkish patients with corneal dystrophies: Five novel variations in CHST6. Molecular Vision 22, 1267–1279 (2016).

12. G. K. Klintworth et al., Macular corneal dystrophy in Saudi Arabia: a study of 56 cases and recognition of a new immunophenotype. Am J Ophthalmol 124, 9–18 (1997).

13. T. K. Bozkurt et al., An 11-Year Review of Keratoplasty in a Tertiary Referral Center in Turkey: Changing Surgical Techniques for Similar Indications. Eye Contact Lens 43, 364–370 (2017).

14. P. Gain et al., Global Survey of Corneal Transplantation and Eye Banking. JAMA Ophthalmol 134, 167–173 (2016).

15. G. K. Klintworth, J. Reed, G. A. Stainer, P. S. Binder, Recurrence of macular corneal dystrophy within grafts. Am J Ophthalmol 95, 60–72 (1983).

16. E. J. C. Alexandre S. Marcon, Christopher J. Rapuano, and Peter R. Laibson, Recurrence of Corneal Stromal Dystrophies After Penetrating Keratoplasty. Cornea 1, 19–21 (2003).

17. J. L. Funderburgh, Keratan sulfate biosynthesis. IUBMB Life 54, 187–194 (2002).

18. T. Saito et al., Sulfation patterns of keratan sulfate in different macular corneal dystrophy immunophenotypes using three different probes. Br J Ophthalmol 92, 1434–1436 (2008).

19. A. H. Plaas et al., Altered fine structures of corneal and skeletal keratan sulfate and chondroitin/dermatan sulfate in macular corneal dystrophy. J Biol Chem 276, 39788–39796 (2001).

20. B. P. Palka et al., Structural collagen alterations in macular corneal dystrophy occur mainly in the posterior stroma. Curr Eye Res 35, 580–586 (2010).

21. S. Murab, S. Chameettachal, S. Ghosh, Establishment of an in vitro monolayer model of macular corneal dystrophy. Lab Invest 96, 1311–1326 (2016).

22. A. T. Hayashida Y, Beecher N, Lewis P, Young RD, Meek KM, Kerr B, Hughes CE, Caterson B, Tanigami A, Nakayama J, Fukada MN, Tano Y, Nishida K, Quantock AJ., Matrix morphogenesis in cornea is mediated by the modification of keratan sulfate by GlcNAc 6-O-sulfotransferase. Proc Natl Acad Sci U S A 36, 13333–13338 (2006).

23. X. C. Zhao et al., The zebrafish cornea: structure and development. Invest Ophthalmol Vis Sci 47, 4341–4348 (2006).

24. D. Puzzolo et al., Structural, ultrastructural, and morphometric study of the zebrafish ocular surface: a model for human corneal diseases? Curr Eye Res 43, 175–185 (2018).

25. A. R. Souza et al., Chondroitin sulfate and keratan sulfate are the major glycosaminoglycans present in the adult zebrafish Danio rerio (Chordata-Cyprinidae). Glycoconj J 24, 521–530 (2007).

26. B. Thisse, C. Thisse (2004) Fast Release Clones: A High Throughput Expression Analysis. ZFIN Direct Data Submission.

27. J. Zhang et al., A comprehensive evaluation of 181 reported CHST6 variants in patients with macular corneal dystrophy. Aging 11, 1019–1029 (2019).

28. M. A. Moreno-Mateos et al., CRISPRscan: designing highly efficient sgRNAs for CRISPR-Cas9 targeting in vivo. Nat Methods 12, 982–988 (2015).

29. C. Kuscu, S. Arslan, R. Singh, J. Thorpe, M. Adli, Genome-wide analysis reveals characteristics of off-target sites bound by the Cas9 endonuclease. Nat Biotechnol 32, 677–683 (2014).

30. K. Kraft et al., Deletions, Inversions, Duplications: Engineering of Structural Variants using CRISPR/Cas in Mice. Cell Rep 10, 833–839 (2015).

31. G. J. Rauch et al. (2003) Submission and Curation of Gene Expression Data. (ZFIN, http://zfin.org).

32. G. K. Klintworth et al., Macular corneal dystrophy. Lack of keratan sulfate in serum and cornea. Ophthalmic Paediatr Genet 7, 139–143 (1986).

33. C. Cursiefen et al., Immunohistochemical classification of primary and recurrent macular corneal dystrophy in Germany: subclassification of immunophenotype I A using a novel keratan sulfate antibody. Exp Eye Res 73, 593–600 (2001).

34. S. Aggarwal, T. Peck, J. Golen, Z. A. Karcioglu, Macular corneal dystrophy: A review. Surv Ophthalmol 63, 609–617 (2018).

35. M. Sabir, The morphogenesis of granular and lattice corneal dystrophy - A mutation combination hypothesis. Med Hypotheses 145, 110291 (2020).

36. D. Xiao et al., Lumican promotes joint fibrosis through TGF-beta signaling. FEBS Open Bio 10, 2478–2488 (2020).

37. S. Brézillon, K. Pietraszek, F. X. Maquart, Y. Wegrowski, Lumican effects in the control of tumour progression and their links with metalloproteinases and integrins. The FEBS Journal 280, 2369–2381 (2013).

38. D. Szklarczyk et al., STRING v10: protein-protein interaction networks, integrated over the tree of life. Nucleic Acids Res 43, D447–452 (2015).

39. C. Thisse, B. Thisse, High-resolution in situ hybridization to whole-mount zebrafish embryos. Nat Protoc 3, 59–69 (2008).

40. C. E. Vejnar, M. A. Moreno-Mateos, D. Cifuentes, A. A. Bazzini, A. J. Giraldez, Optimized CRISPR-Cas9 System for Genome Editing in Zebrafish. Cold Spring Harb Protoc 2016 (2016).

41. L. E. Jao, S. R. Wente, W. Chen, Efficient multiplex biallelic zebrafish genome editing using a CRISPR nuclease system. Proc Natl Acad Sci U S A 110, 13904–13909 (2013).

