## Supplementary Information for "Zebrafish *carbohydrate sulfotransferase 6* (*chst6*) mutants provide a preclinical model for macular corneal dystrophy"

**Methods**

**Verification of Mutagenesis and Genotyping**

For obtaining DNA lysates embryos (up to 8), or fin clips from individual fish were lysed in 50 μl of 50 mM NaOH. incubating at 95 °C for 30 min, and finally neutralization of the solution with 50 μl of 100 mM Tris-HCl (pH:8.0). For evaluating the efficiency in mosaic F_0_ pools of 8 embryos were used, while Founder Fish (FF) screening was done on at least 8 individual F_1_ embryos, for each embryo 5 μl NaOH and 5 μl Tris-HCl was used. For adult identification fin clip DNA was used. Genomic DNA was amplified with Forward 5’CCACTGCTAATGTCGTGATGTT3’ and Reverse 3’CGCAGTGCTTTTTGCAGTCT5’ primers. Deletion mutants were identified by shift of mobility on agarose gel. Small indel mutations were identified by T7 Endonuclease (NEB, MO302L) mismatch assay, according to product manual.

For sequence analysis genomic DNA was amplified with Forward:5’CCACTGCTAATGTCGTGATGTT3’, Reverse 3’CGCAGTGCTTTTTGCAGTCT5’ primers and Sanger sequencing was done. Sequence analysis was performed with the Clustal Omega web-based software and SnapGene software.

**Histopathology protocols**

Immunofluorescence staining of adult cornea was performed on cryosections. One eye of euthanized zebrafish was surgically removed and fixed in %4 formaldehyde, overnight at room temperature, rest of the fish body was used for paraffin embedding. Fixed eye tissue was rinsed with PBS and immersed in 30% sucrose for 3 days at 4°C until eyes sank to the bottom of the tube. Next, tissue was transferred into OCT (Sakura, Tissue-Tek, 4583), kept at -20°C for 30 min and -80°C overnight. 14 µm sections were taken with a cryostat (Leica, CM 1950), and transferred onto slides (Epredia, J1800AMNZ). Slides were stored at -80°C, until further processing. IF staining slides were rehydrated with 1X PBDT (0.1% Tween-20, %1 DMSO in PBS) at room temperature for 5 min. Slides were blocked with blocking solution (5% normal goat serum in PBDT) for 1-2 hours at RT. Then samples were incubated with primary antibody overnight at 4°C in a humidified chamber, rinsed with 1X PDT 3 times, incubated with secondary antibody Goat Anti-Mouse, Alexa Fluor 594 (1:400, Invitrogen, A11005) for 2 hours at room temperature. DAPI (Sigma, D9542-5MG) was used for nuclear staining. 40X Water and 63X objectives were used to capture Z-stacks, background subtraction and maximum Z-projection were applied with ImageJ.

Alcian blue staining was performed on paraffin sections of whole body. Euthanized zebrafish were fixed in 10% neutral buffered formalin for at least six days. Following fixation, the fish were treated with a 0.35 M EDTA/PBS solution for five days to soften the bones. Tissue was passed through ethanol series (70%, 80%, 96%, 2 times 100%) to dehydrate. Samples were incubated in fresh xylene 2 times, and in paraffin at 67°C. After processing whole body or heads are embedded in paraffin using base mold, blocks were frozen at -20 °C for at least 2-3 hr. 5 µm thick sections were placed on poly-L-lysine coated slides (Laborant, ANT-MS90PS01W). Alcian blue/PAS staining (Bio-Optica, 04-163802) was performed according to manufacturer directions.

**Results**

**Success rate of each strategy**

gRNA injected fish embryo clutches were raised after confirmation of mutagenesis occurrence in siblings by mosaic embryo genomic PCR followed by gel mobility shift for deletion mutations or T7EI assay for indel mutations. The mosaic adults were screened with same gel analysis method in offspring F_1_ embryos. The number of mosaic adults that were screned, number of candidate founder fish indicated by positive result in gel analysis, number of confirmed fonder fish by sequencing are reported in Table S1.

**Table S1. Statistics of gRNA success rate**

| **gRNA(s)** | **# of Mosaic Adults (F_0_)** | **# of Founder Fish canditate**  **(Gel analysis)** | **# of Founder Fish**  **Confirmed with sequencing** | **% Positive** |
| --- | --- | --- | --- | --- |
| gRNA5 | 95 | 6 | 0 | %0 |
| gRNA3+5 | 20 | 1 | 1 | % 5 |
| gRNA1 | 45 | 13 | 13 | % 28.8 |
| gRNA1+2 | 135 | 6 | 6 | % 4.4 |

**Failed attempt to delete the coding sequence**

In an attempt to generate coding sequence deletion mutation mixture of gRNA3, gRNA5 and Cas9 mRNA was injected into 1-cell stage embryos. High rate of embryonic lethality was observed when *nlsCas9nls* mRNA was used. In order to limit Cas9 activity to primordial germ cells *nanos_nlsCas9nls*, which partially decreased embryonic lethality, and a total of 20 adult fish were obtained (Fig. S1). When genomic sequence was amplified by PCR WT (1.3 kb) and mutant (350 bp) bands were detected in mosaic adults (Fig. S1B). However a 150-160 bp mutant DNA band was expected if the targeted DSBs were to occur. Presence of A large deletion was detected in 1 fish (FF5) (FigS2-B). In the DNA analysis of the single embryo (Fig. S2D), a DNA band of approximately 350 bp was seen and F_1_ generation was obtained from this fish. According to this analysis, only a mutant DNA band of approximately 350 bp in length is seen in fish 5.1 and 5.3. When DNA sequence analysis was performed from the F_1_ heterozygous generation of mutant fish obtained, it was seen that a 943 bp region, nucleotides 236-1179 of the coding sequence were deleted was deleted. Alignment analysis showed that no mutation occurred in the region targeted with gRNA5, but deletion started 200 bp below this region. While gRNA3 guided DSB occurred in the predicted location. According to the alignment of protein sequence (Fig. S1 F) although a part of the gene was deleted in these mutants, the PAPS region remained the same.

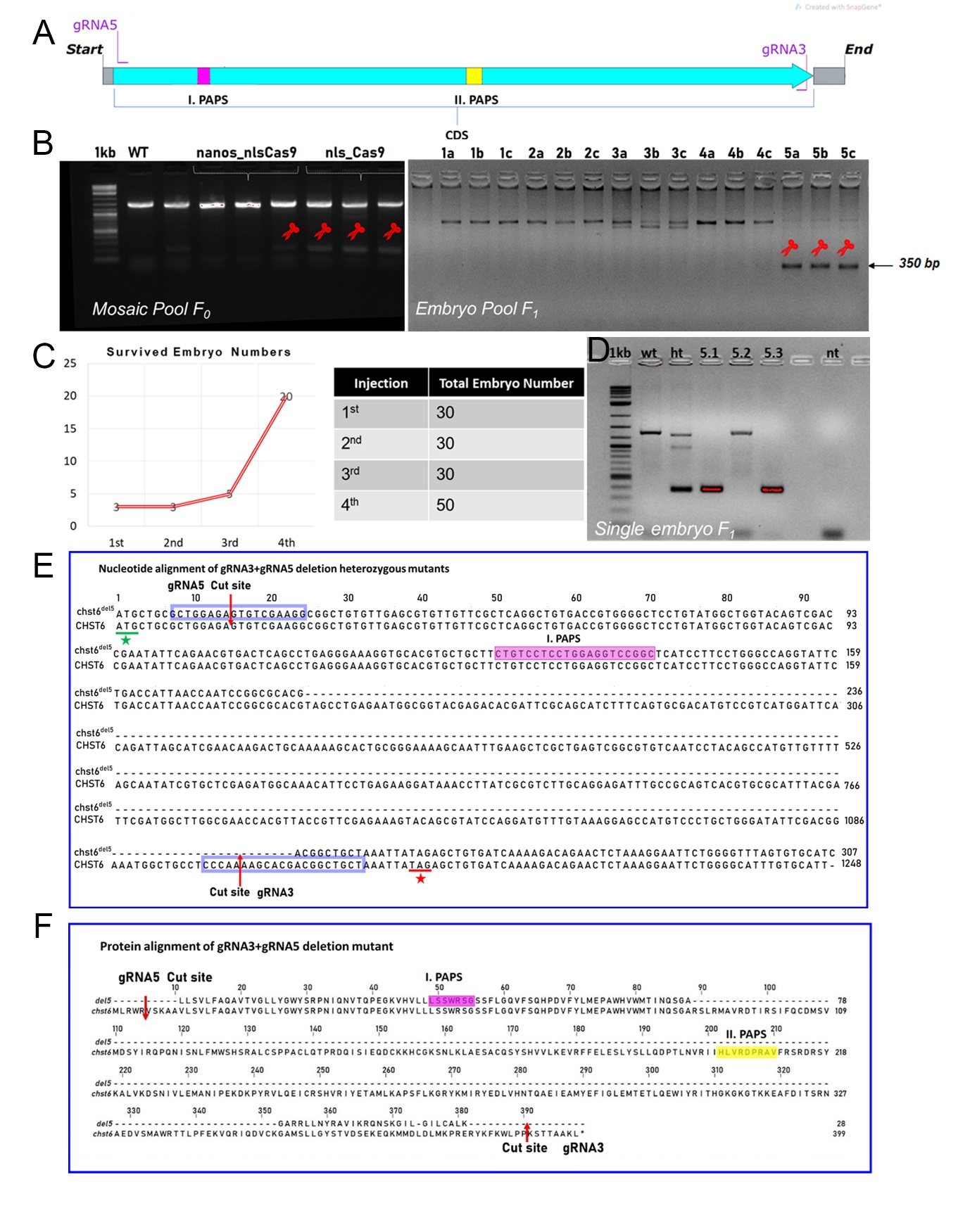

**Figure S1: Deletion of the entire coding sequence of *chst6* was not possible**  **A)** Graphical representation of *chst6* gene cds and gRNA binding sites **B)** DNA gel electrophoresis results showing wt and mutant DNA amlified by PCR. (Left) Pools of injected embryos were analysed and the clutched in which mutant DNA band (red scisoors) were raised. (Right) Progeny of mosaic adults were screened for mutagenesis in 3 pools of 8 embryos. Image shows a 350 bp mutant DNA fragment in progeny of adult no 5, whereas only a small deletion was deteted in progeny of adult no 3. **C)** Number of adults that survived after 4 rounds of injection is shown, total number of embryos raised are shown in the table. nanos_nlsCas9nls was used in 4^th^ injection to prevent lethality. **D)** Mutant DNA analysis of single embryos obtained from adult fish no 5. Sequence of del5 mutant obtained from FF5 is shown **E)** as nucleotide alignment and **F)** protein alignment. 1^st^ PAPS is shown in pink, and 2^nd^ PAPS is shown in yellow.

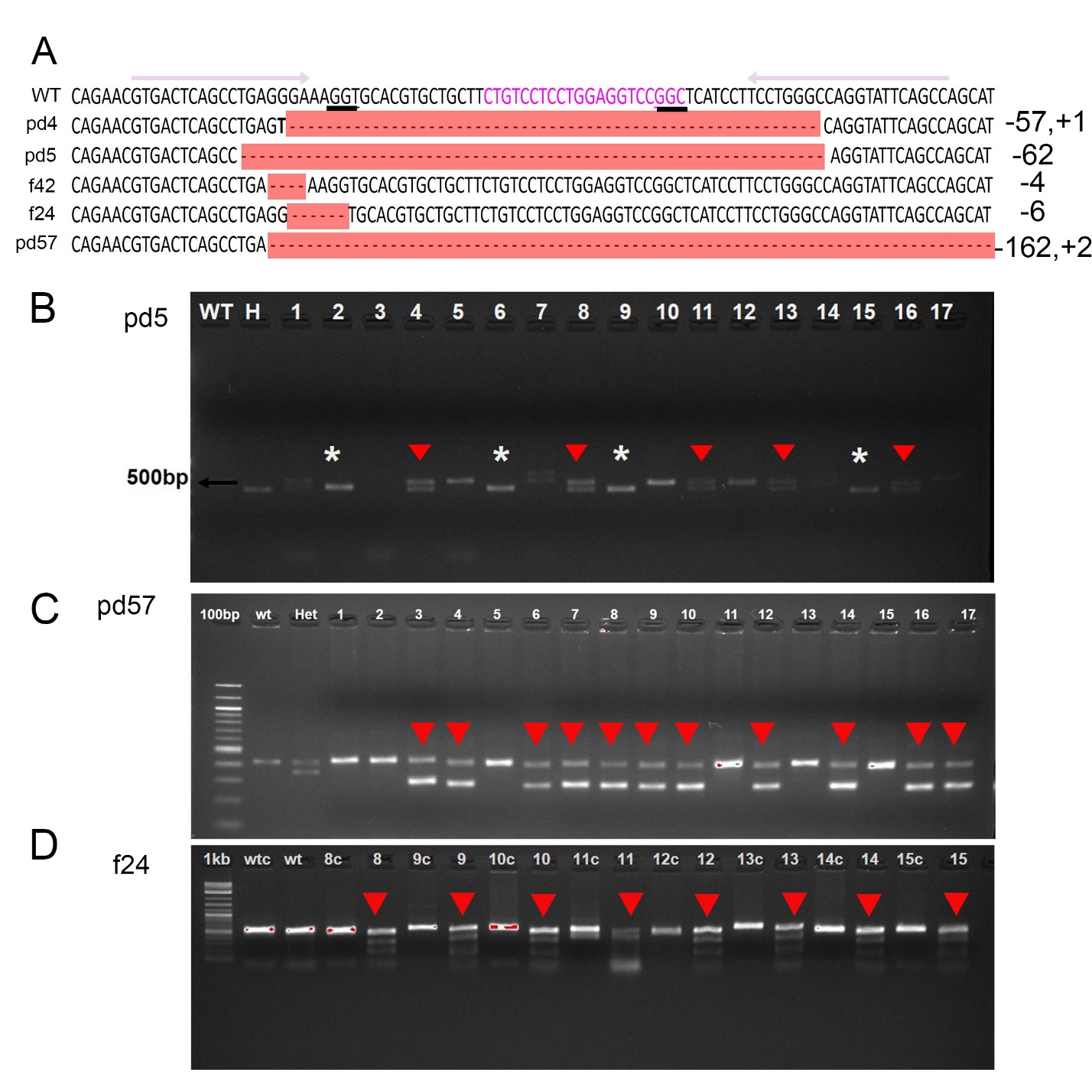

**Figure S2** : **A)** Multiple sequence alignment of chst6 mutant alleles. chst6^f24^ has a 6 nucleotide deletion whereas chst6^f42^ has 4 nucleotide deletion. chst6^pd4^ and chst6^pd5^ has small deletions od 57 and 62 bp, respectively.Wheras chst6^pd57^  has a larger deletion of 162 bp. Pink arrow shows gRNAs and PAM sites are underlined black. Nucleotide coding for 1^st^ PAPS are colored magenta. **B-D)** Representative images of mutagenesis screening gel analyses. **B)** F_1_ adults raised from founder fish 5 were genotyped from fin clip DNA. *chst6^pd5/+^* heterozygous mutants had a double band pattern. **C)** F_1_ adults raised from founder fish 57 were genotyped from fin clip DNA. *chst6^pd57/+^* heterozygous mutants had a double band pattern. **D)** Heterozygous mutants carrying the chst6^f24/+^ allele were identified with T7EI analysis control reaction and digest rection were loaded consecutively, heterozygous mutants were positive for T7EI digest. Heterozygous mutants are indicated with red arrowheads.

**Table S2. Table of** **the number of heterozygous and homozygous fish of selected alleles.**

| **Allele Name** | **Initial number of fish** | **Number of total**  **F_2_ progeny** | **Number of F_2_ heterozygous embryos** | **Number of F_2_ homozygous embryos** |
| --- | --- | --- | --- | --- |
| chst6^pd4/4?^ | 40 | 34 | 15 | 7 |
| chst6^pd5/5?^ | 50 | 26 | 19 | 3 |
| chst6^f42/42?^ | 50 | 36 | 15 | 10 |

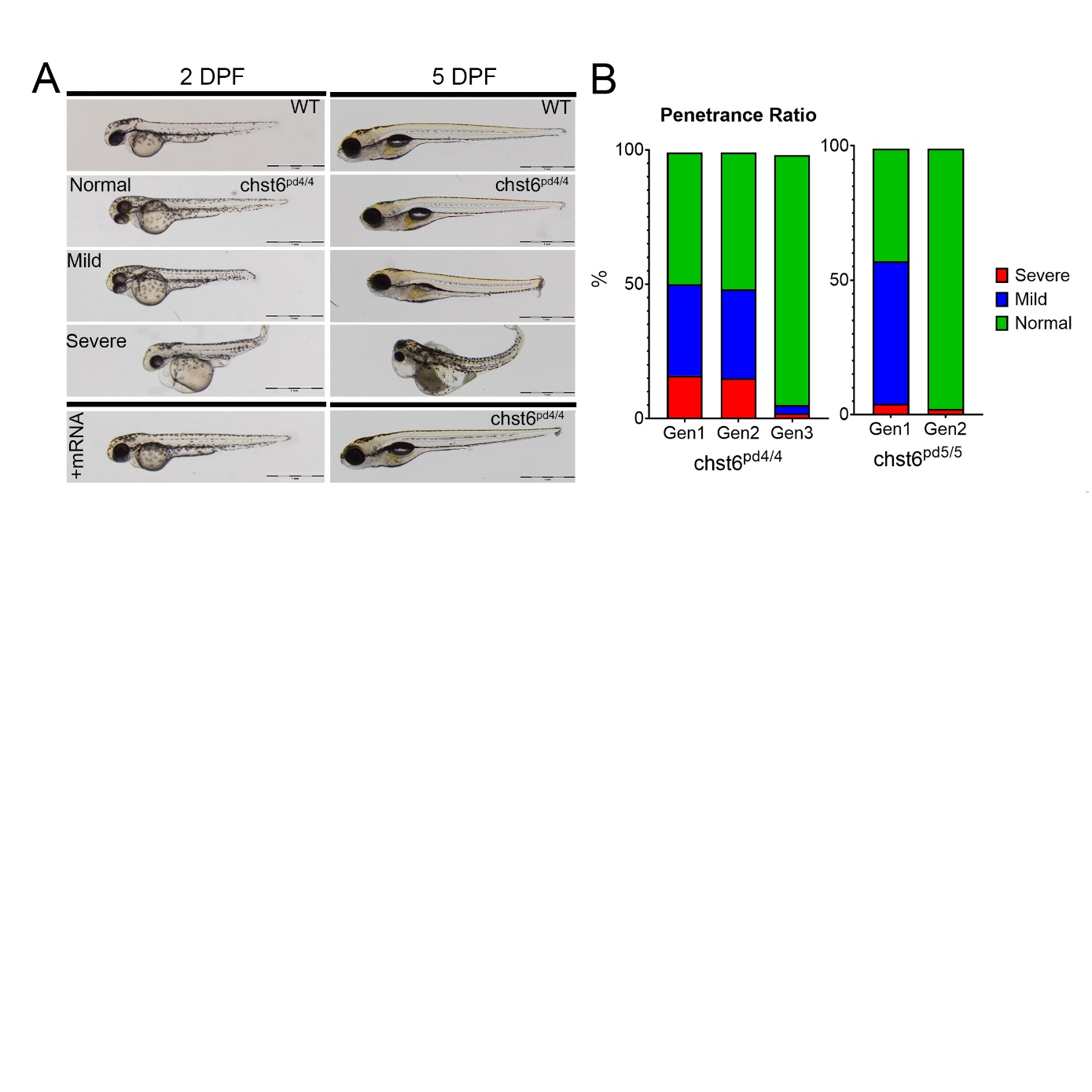

**Figure S3**. **Morphological defects were observed in *chst6* mutant zebrafish larvae**

**A)** Homozygous mutatns displayed morphlogical defects at 2 dpf and 5 dpf, and the severe ones did not survive. The defects were fully rescued when wt chst6 mRNA was injected to 1 cell-stage embryo. **B)** The penetrance of morphological defect phenotype is shown for *chst6^pd4/4^* and *chst6^pd5/5^* alleles in different generations. The defects were oserved in less than 5% of the mutants in generation (gen) 2 and gen 3.

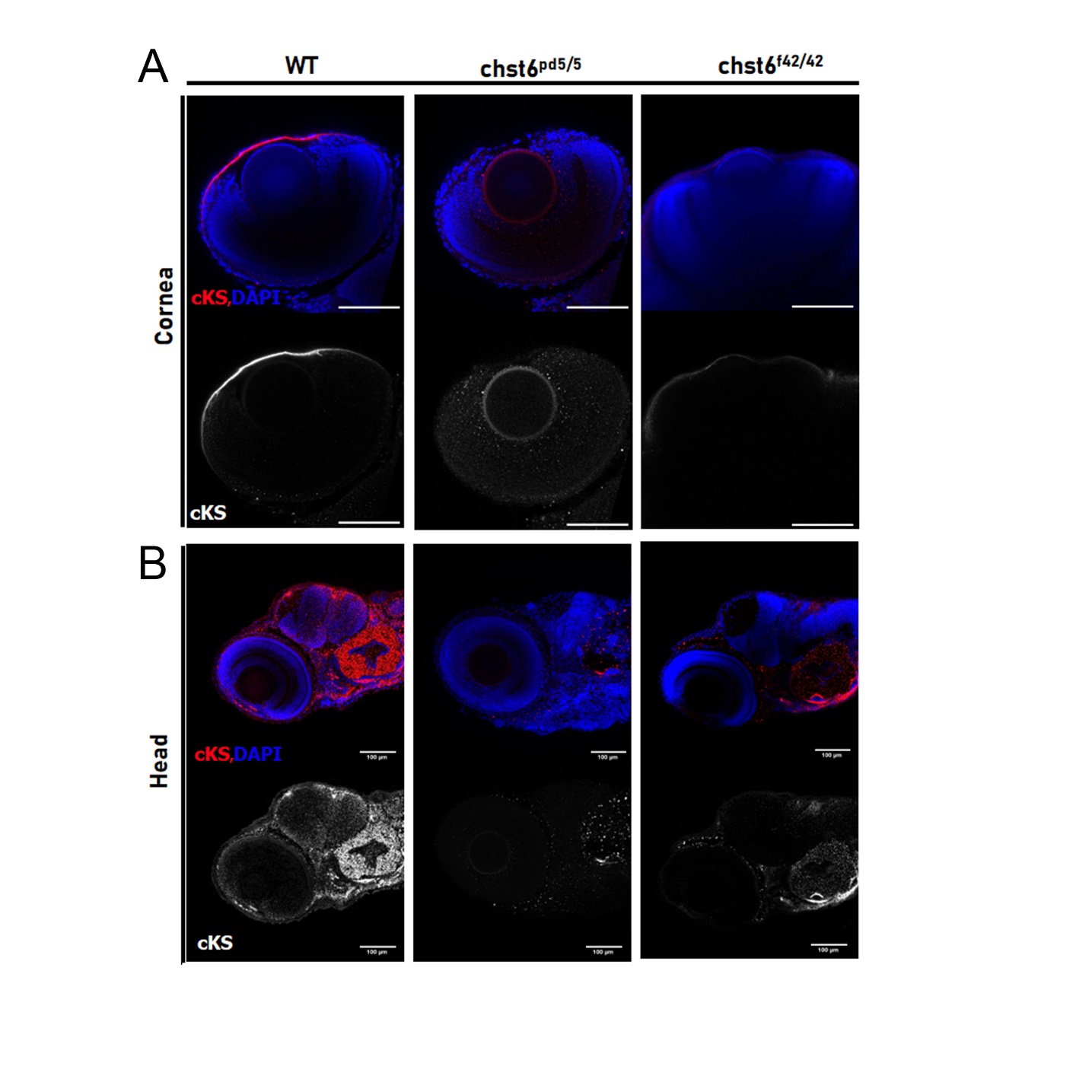

**Figure S4. RZebrafish mutants have defective keratan sulfate sulfation**

Representative images of cKS antibody staining: **A)** eye of wt, chst6^pd5/pd5^ and chst6^f42/f42^ mutant larva.

**B)** head of wt, chst6^pd5/pd5^ and chst6^f42/f42^  mutant larva. cKS: red, DAPI: blue, Scale bar: 100 µm

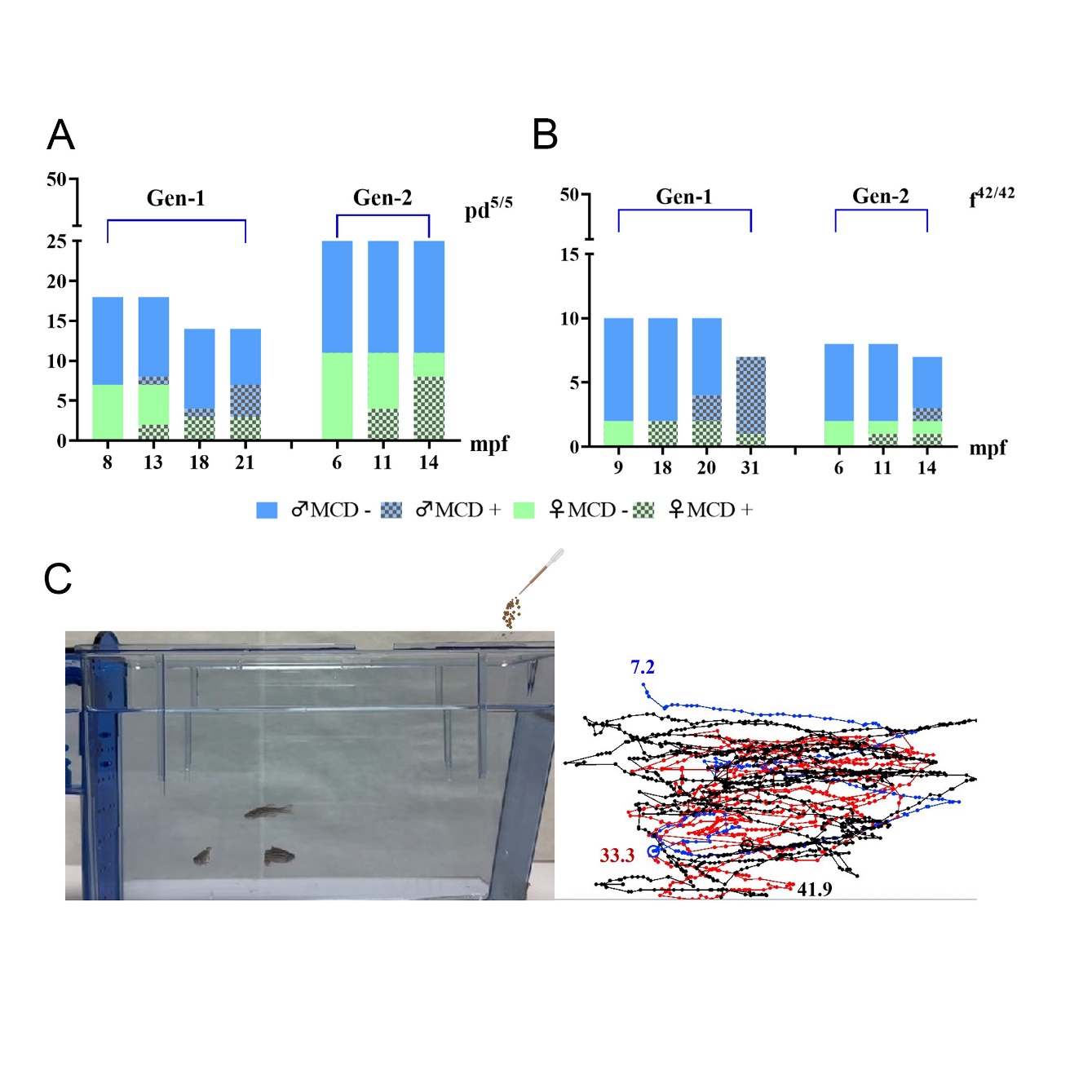

**Figure S5 :** Graphics showing the total number of adults and incidence of opacity in the eyes over time. Green bar: females, blue bar: males, dashed bars indicate occurrence of opaque aggregates. **A)** chst6^pd5/5^ **B)** chst6^f42/42^ mutants in geeration 1 and 2. **C)** Vision test was performed on chst6^pd4/4^ 23 mpf adult fish Swim path and duration of fish were tracked. Track was ended if the fish reached food and time stamp is displayed at the end of each track. The mutants swam for 33.3 and 41.9 seconds without noticing the food.

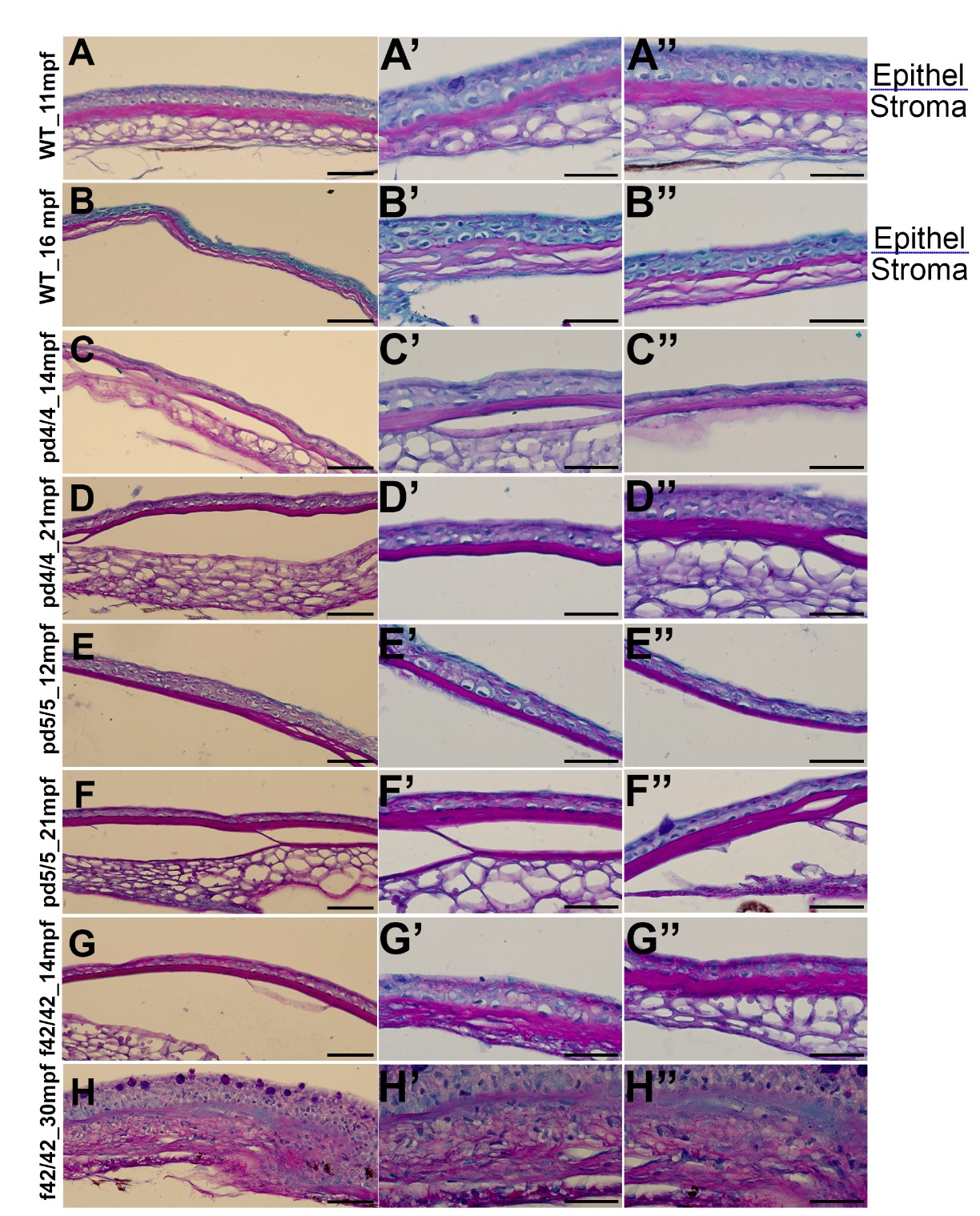

**Figure S6. Figure 5: Zebrafish with MCD have alcian blue positive aggregates in the cornea**

Alcian blue/PAS-stained cornea tissue of **A)** 11 mpf WT, **B)** 16 mpf WT, **C)** 14 mpf *chst6^pd4/pd4^* , **D)** 21 mpf *chst6^pd4/^pd4*, **E)** 12 mpf *chst6^pd5/pd5^*, **F)** 21 mpf *chst6^pd5/pd5^* , **G)** *14 mpf chst6^f42/42^* **H)** 30 mpf *chst6^f42/42^* are shown. Scale bar 50 µm. Zoom images recorded with 100X objective are shown in **A’-A’’)** 11 mpf WT, **B’-B’’)** 16 mpf WT, **C’-C’’)** 14 mpf *chst6^pd4/pd4^* , **D’-D’’)** 21 mpf *chst6^pd4/^pd4*, **E’-E’’)** 12 mpf *chst6^pd5/pd5^*, **F’-F’’)** 21 mpf *chst6^pd5/pd5^* , **G’-G’’)** *14 mpf chst6^f42/42^* **H’-H’’)** 30 mpf *chst6^f42/42^* 20C overall eye structure. Scale bar: 30 µm.

**Supplementary videos:**

**SuppVideo1:** wt feeding video

**SuppVideo 2:** mutant (*chst6^pd4/pd4^*) feeding video

**SuppVideo 3:** mutant (*chst6^f42/f42^*) feeding video
